# Basal mitophagy is widespread in *Drosophila* but minimally affected by loss of Pink1 or parkin

**DOI:** 10.1101/235077

**Authors:** Juliette J. Lee, Alvaro Sanchez-Martinez, Aitor Martinez Zarate, Cristiane Benincá, Ugo Mayor, Michael J. Clague, Alexander J. Whitworth

## Abstract

Parkinson’s disease factors, PINK1 and parkin, are strongly implicated in stress-induced mitophagy in vitro, but little is known about their impact on basal mitophagy *in vivo*. We generated transgenic *Drosophila* expressing fluorescent mitophagy reporters to evaluate the impact of *Pink1/parkin* mutations on basal mitophagy under physiological conditions. We find that mitophagy is readily detectable and abundant in many tissues including Parkinson’s disease relevant dopaminergic neurons. However, we did not detect mitolysosomes in flight muscle. Surprisingly, in *Pink1* or *parkin* null flies we did not observe any substantial impact on basal mitophagy. As these flies exhibit locomotor defects and dopaminergic neuron loss, our findings raise questions about current assumptions of the pathogenic mechanism associated with the PINK1/Parkin pathway. Our findings provide evidence that Pink1 and parkin are not essential for bulk basal mitophagy in *Drosophila*. They also emphasize that mechanisms underpinning basal mitophagy remain largely obscure.

**Summary:** PINK1/parkin are key mediators of stress-induced mitophagy *in vitro* but their impact on basal mitophagy *in vivo* is unclear. Novel *Drosophila* reporters lines reveal abundant mitophagy in many tissues including dopaminergic neurons but is unaffected by loss of PINK1/parkin.

## Introduction

Mitochondria are essential organelles that perform many critical metabolic functions but are also a major source of damaging reactive oxygen species (ROS) and harbor pro-apoptotic factors. Multiple homeostatic processes operate to maintain mitochondrial integrity, however, terminally damaged organelles are degraded through the process of targeted mitochondrial autophagy (mitophagy) to prevent potentially catastrophic consequences. Such homeostatic mechanisms are particularly important for post-mitotic, energetically demanding tissues such as nerves and muscles. Two proteins linked to Parkinson’s disease (PD), Parkin, a cytosolic ubiquitin E3 ligase, and PINK1, a mitochondrially targeted kinase, have been shown to play key roles in mitophagy.

PINK1-Parkin mediated mitophagy has been intensively investigated and many of the molecular mechanisms have been elucidated (reviewed in (Pickrell and Youle, 2015; Yamano et al., 2016). Briefly, upon loss of mitochondrial membrane potential as occurs in damaged or dysfunctional mitochondria, PINK1 accumulates on the outer mitochondrial membrane (OMM) and initiates the mitophagy signal by phosphorylating both ubiquitin and Parkin, which promotes its activity depositing more ubiquitin for subsequent phosphorylation. This process acts as a feedforward mechanism ultimately decorating the OMM with phospho-ubiquitin chains that are then recognized by ubiquitin adaptor proteins leading to the engulfment of the depolarized mitochondria by autophagosomes.

As a pathogenic mechanism, failure in mitophagy offers an attractive explanation for multiple, long standing observations that implicate mitochondrial defects in the pathogenicity of PD, such as systemic mitochondrial complex I deficits and high levels of mtDNA mutations in PD patients (Bender et al., 2006; Dias et al., 2013; Exner et al., 2012; Kraytsberg et al., 2006; Schapira et al., 1989). In addition, the unique physiological characteristics of *substantia nigra* neurons, such as their extensive arborization, myriad synaptic connections and continuous pacemaking activity, place an extreme demand on mitochondrial function to meet their high energy demand and calcium buffering requirements, which may explain the selective vulnerability of these neurons to loss of mitochondrial integrity (Sulzer and Surmeier, 2013).

Many of the molecular details of PINK1/Parkin-induced mitophagy have been elaborated in cultured cells that over-express Parkin following acute mitochondrial depolarization or toxification (Pickrell and Youle, 2015; Yamano et al., 2016). However, relatively little is known about mitophagy under physiological conditions *in vivo* (Cummins and Gotz, 2017; Rodger et al., 2017; Whitworth and Pallanck, 2017). One of the limitations to studying mitophagy *in vivo* has been the paucity of suitable reporters. Recently, two *in vivo* mitophagy reporter models have been described in mice (McWilliams et al., 2016; Sun et al., 2015). One uses a mitochondrial matrix-targeted pH-sensitive variant of GFP (mt-Keima) while the other uses a tandem GFP-mCherry fusion protein targeted to the OMM (called “mito-QC”). Both systems exploit pH-sensitive properties of mKeima and GFP respectively to enable the differential labelling of mitochondria in the acidic microenvironment of the lysosome as a proxy end-point readout. Initial reports on these two reporter lines have revealed a surprisingly widespread and heterogeneous distribution of basal mitophagy, however, the involvement of PINK1 and Parkin has not been addressed yet.

We have generated *Drosophila* lines expressing these mitophagy reporters to provide the first global view of the prevalence of mitophagy across the organism and to determine the relative contribution of Pink1 and parkin to basal mitophagy. Analyzing mitophagy in *Drosophila* offers a valuable opportunity to interrogate the physiological requirement of Pink1-Parkin in this process *in vivo*. In contrast to mammalian models, loss of *Drosophila Pink1* or *parkin* leads to robust phenotypes in locomotor activity and dopaminergic (DA) neuron loss (Clark et al., 2006; Greene et al., 2003; Park et al., 2006), indicating their critical function in the related neuromuscular tissues. We find that while basal mitophagy is widespread in many tissues in *Drosophila*, the incidence of mitolysosomes, and hence, basal mitophagy, is unaffected by loss of Pink1 or parkin. Moreover, flight muscle, which shows the most robust disruption upon *Pink1/parkin* mutation (Clark et al., 2006; Greene et al., 2005; Greene et al., 2003; Park et al., 2006), does not reveal a detectable mitophagy signal. Hence, we propose that Pink1 and parkin are largely dispensable for basal mitophagy have mitophagy-independent functions in neuromuscular tissues in *Drosophila*.

## Results and discussion

### Validation of mitophagy reporters in *Drosophila*

We first generated GAL4/UAS-inducible transgenic lines to express either mito-QC or mt-Keima. Multiple lines were established and immunoblotting confirmed comparable expression between lines (Figure S1A-B). Expression of these reporters in all lines appeared benign and did not noticeably affect development or viability. To ensure that the expression of these reporters did not overtly interfere with normal cellular functions *in vivo*, particularly in the sensitive neuromuscular systems of interest here, we analyzed locomotor activity in flies expressing high levels of the mito-QC or mt-Keima in all tissues and observed no significant impact (Figure S1C-F).

Previous work has demonstrated the appropriate targeting of mito-QC and mt-Keima to mitochondria *in vitro* and *in vivo*, and that ‘spectral shifted’ puncta of more acidic conditions indeed co-localized with lysosomes. Nevertheless, we also sought to verify these conditions in our *Drosophila* lines. As expected, we observed substantial co-localization of mito-QC and mt-Keima with the mitochondrial protein ATP5A in two tissues highly amenable for mitochondrial imaging analysis, larval epidermal cells and adult flight muscle (Figure S2A-C). The ‘spectral shift’ that marks mitolysosomes of mito-QC occurs by the selective quenching of GFP but not mCherry, resulting in ‘red-only’ puncta (Figure S2D). Similarly, for mt-Keima the fluorescence spectra shifts to reflect an increased signal from excitation at 561 nm when under more acidic conditions (Figure S2E). With both mitophagy reporters we observed a striking co-localization of ‘mitophagy’ puncta with LysoTracker (Figure S2D, E).

To further validate these puncta as mitolysosomes we assessed their formation in tissue lacking the canonical autophagy pathway generated by knockdown of the key autophagy factor *Atg5*. Mitolysosome signal reported by both mito-QC and mt-Keima were almost entirely absent in *Atg5*-RNAi epidermal cells (Figure S3A), consistent with their appearance reflecting mitophagy. Moreover, it was previously reported that iron chelation by deferiprone induced mitophagy reported by mito-QC (Allen et al., 2013). We also observed an increase in mitolysosomes reported by mito-QC and mt-Keima by feeding animals deferprione (Figure S3B). Taken together these findings validate the ‘mitophagy’ signal of mito-QC and mt-Keima as faithful reporters of mitolysosomes in *Drosophila*, similarly to previously characterized mouse models (McWilliams et al., 2016; Sun et al., 2015).

### Mitophagy is widespread in multiple *Drosophila* tissues

We next determined the prevalence of mitolysosomes under basal conditions in multiple tissues. In larvae we analyzed epidermal cells, since they have an elaborate mitochondrial morphology, and the ventral ganglion, a major portion of the central nervous system (CNS). Here, we directly compared the mitolysosome signal reported by mito-QC and mt-Keima. Microscopic analysis revealed that mitolysosomes are abundant under basal conditions in both epidermal cells and CNS (Figure 1). In contrast, mitolysosomes were almost undetectable in larval body wall muscles (data not shown). Importantly, mito-QC and mt-Keima revealed a very similar abundance and distribution of mitolysosomes substantiating their utility as equivalent mitophagy reporters.

**Figure 1.**
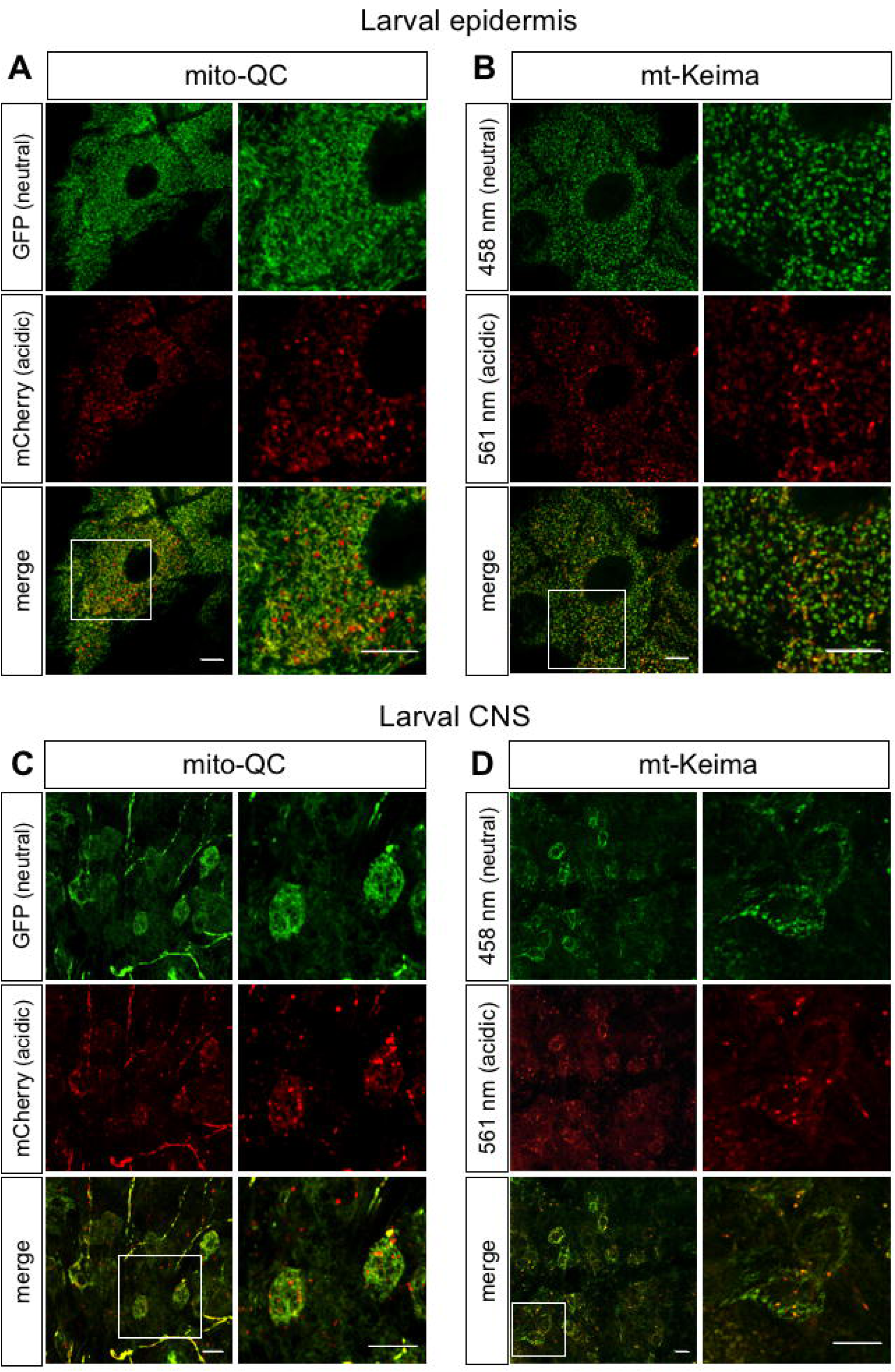
Comparison of mito-QC and mt-Keima mitophagy reporters in larval tissues of *Drosophila*. Confocal microscopy analysis of (A, B) larval epidermal cells and (C, D) the ventral ganglion of the central nervous system (CNS), visualizing mito-QC and mt-Keima as indicated. Expression was induced by ubiquitous GAL4 drivers. Genotypes analyzed were (A, C) *da-GAL4/UAS-mito-QC*, and (B, D) *tub-GAL4*, *UAS-mt-Keima*. Scale bars = 10 μm.

During this stage of analysis we found that the mt-Keima signal, which was already markedly weaker than that generated by mito-QC, was rapidly bleached upon extended exposure or repeated scanning. Repeated imaging was necessary to achieve adequate penetration into complex tissue and for z-stack (3D) reconstruction for mitolysosome quantification. For this reason the subsequent analyses mainly focused on analyzing mito-QC.

We next analyzed adult tissues where post-mitotic, highly energetic tissues would be expected to accumulate more mitochondrial damage and therefore likely undergo more mitophagy. Again, we observed abundant mitolysosomes widespread in the adult brain (Figure 2). PD is associated with the preferential degeneration of dopaminergic (DA) neurons, which is reproduced in multiple *Drosophila* models of PD, including by cell autonomous loss of *Pink1/parkin* (Hewitt and Whitworth, 2017). Mito-QC was selectively expressed in dopaminergic neurons via the tyrosine hydroxylase (TH) GAL4 driver. Analyzing the PPL1 cluster that has most consistently been shown to be affected in *Pink1/parkin* mutants we observed a substantial mitophagy signal in the DA neurons (Figure 2). The amount of mitophagy signal did not markedly increase with ageing.

**Figure 2.**
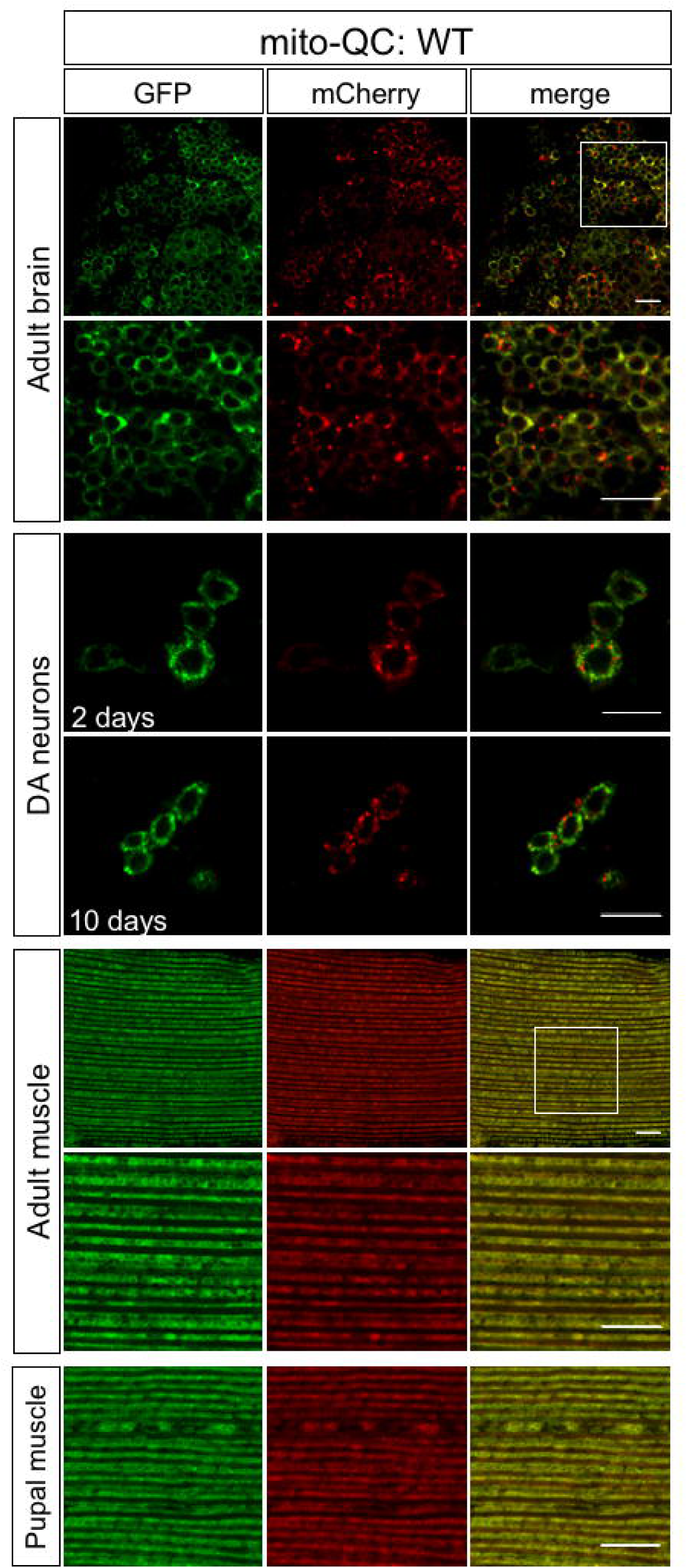
Mitolysosome analysis with mito-QC in adult *Drosophila* tissues. Confocal microscopy analysis of mito-QC reporter in adult brain, dopaminergic (DA) neurons and flight muscle, as indicated. Mitolysosomes are evident as GFP-negative/mCherry-positive (‘red-only’) puncta. Quantification of mitolysosomes are shown in Figures 3 and 4. Genotypes analyzed were (adult brain) *nSyb-GAL4/UAS-mito-QC*, (DA neurons) *TH-GAL4/UAS-mito-QC*, and (adult and pupal muscle) *Mef2-GAL4/UAS-mito-QC*. Scale bars = 10 μm.

**Figure 3.**
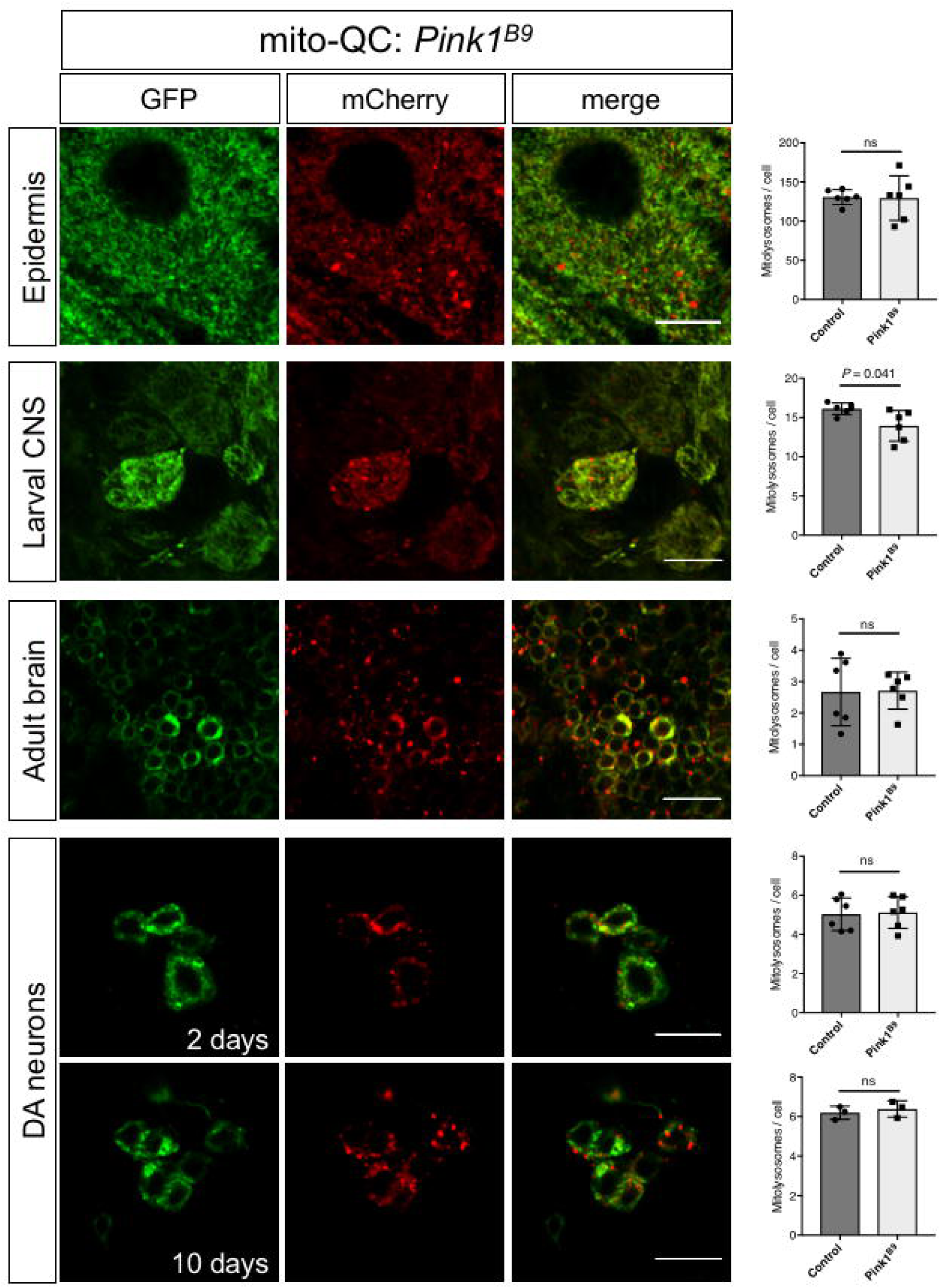
Basal mitophagy is minimally affected in *Drosophila Pink1* mutants. Confocal microscopy analysis of mito-QC reporter in larval epidermis and CNS, adult brain, dopaminergic (DA) neurons and flight muscle, as well as pre-adult (pupal) flight muscles, as indicated, in *Pink1*^*B9*^ mutant animals. Mitolysosomes are evident as GFP-negative/mCherry-positive (‘red-only’) puncta. Scale bars = 10 μm. Charts show quantification of mitolysosomes (see Methods), and show mean ± SD of n = 6 animals. Quantification of WT control samples is the same as shown in Figure 4. Statistical significance determined by Welch’s *t*-test; ns = non-significant. Genotypes analyzed were (epidermis and larval CNS) *Pink*^*B9*^*/Y;da-GAL4/UAS-mito-QC*, (adult brain) *Pink*^*B9*^*/Y;nSyb-GAL4/UAS-mito-QC*, (DA neurons) *Pink*^*B9*^*/Y;TH-GAL4/UAS-mito-QC*, and (adult muscle) *Pink*^*B9*^*/Y;Mef2-GAL4/UAS-mito-QC*.

**Figure 4.**
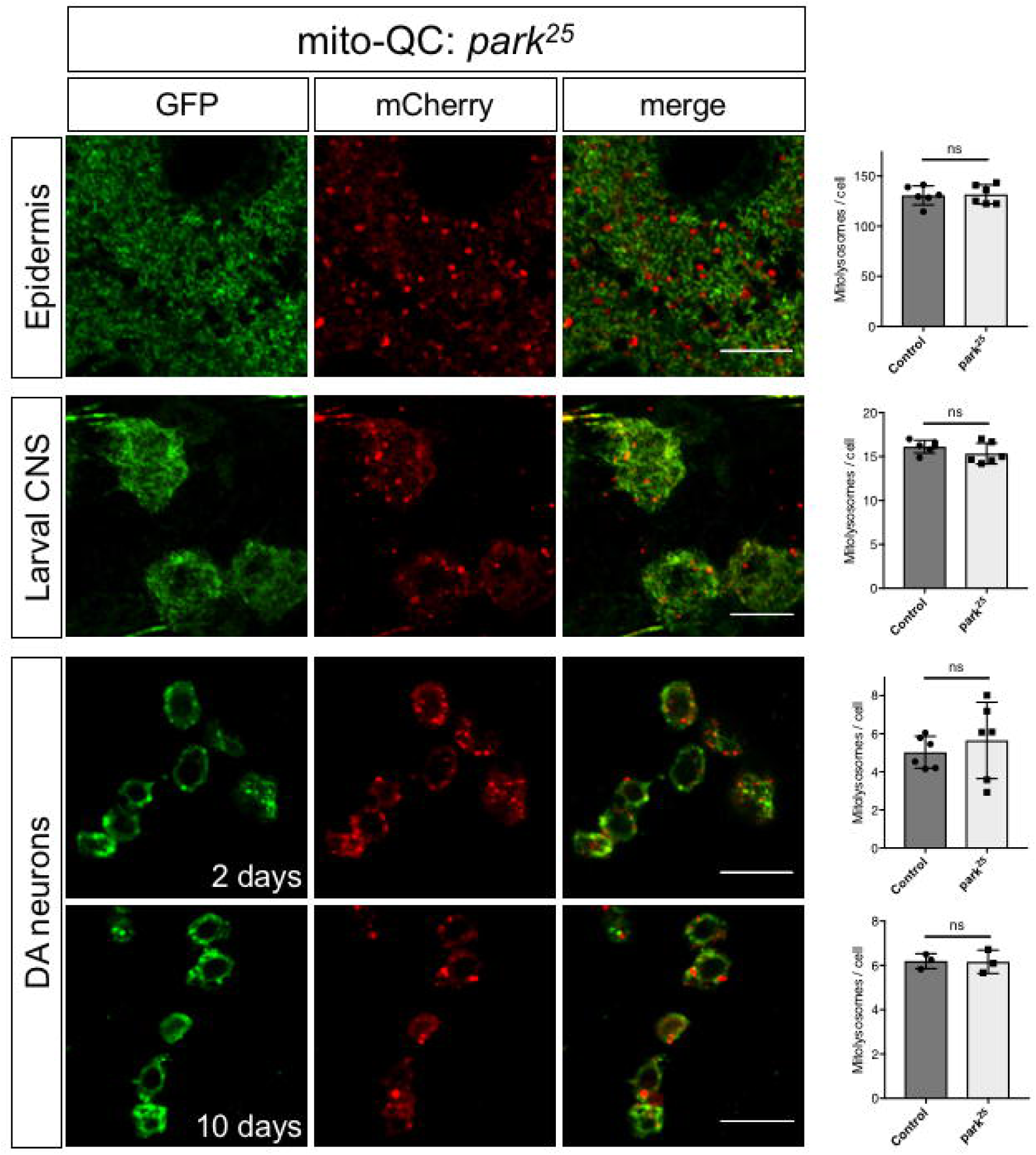
Basal mitophagy is minimally affected in *Drosophila parkin* mutants. Confocal microscopy analysis of mito-QC reporter in larval epidermis and CNS, and adult dopaminergic (DA) neurons, as indicated, in *park*^*25*^ mutant animals. Mitolysosomes are evident as GFP-negative/mCherry-positive (‘red-only’) puncta. Scale bars = 10 μm. Charts show quantification of mitolysosomes (see Methods), and show mean ± SD of n = 6 animals. Quantification of WT control samples is the same as shown in Figure 3. Statistical significance determined by Welch’s *t*-test; ns = non-significant. Genotypes analyzed were (epidermis and larval CNS) *UAS-mito-QC; park*^*25*^, *da-GAL4/park*^*25*^ and (DA neurons) *park*^*25*^, *TH-GAL4/park*^*25*^, *UAS-mito-QC*.

Somewhat surprising, but consistent with observations in larvae, we did not observe any appreciable mitolysosomes in adult muscle tissue (Figure 2). This is notable for two reasons; first, adult flight muscle in particular has extremely abundant mitochondria necessary to power flight. Second, *Pink1* and *parkin* mutant *Drosophila* display robust degeneration of flight muscles. It could be reasonably assumed that this degeneration results from a failure in mitophagy induction. We reasoned that analyzing adult tissue could possibly have missed a critical period when mitophagy is active and required for muscle integrity, such as when the tissue is being formed in development. To address this, we analyzed flight muscle of the pupal stage when disruption of muscle integrity first occurs in *Pink1/parkin* mutants (Clark et al., 2006; Greene et al., 2005). However, we also could not detect mitolysosomes at these earlier time points (Figure 2). Taken together, from these observations we surmise that mitophagy is abundant in various tissues during development and in adult flies, especially in neuronal tissue, but is not detectable in muscle tissue at either stage.

### Pink1 and parkin are largely dispensable for basal mitophagy in *Drosophila*

Mammalian PINK1 and parkin have been described to play important roles in toxin-induced mitophagy, however, the impact of PINK1 and parkin on basal mitophagy *in vivo* has not been reported. To address this, we combined our mitophagy reporters with well-characterized *Pink1* and *parkin* mutants. Analyzing the previously described tissues where mitolysosomes were abundant, we found that mitophagy was not dramatically reduced in genetic null mutants for *Pink1* (Figure 3) and *parkin* (Figure 4). To quantify the degree of mitophagy we applied a semi-automated quantification analysis to segment 3-dimensional images to identify and quantify *bone fide* mitolysosomes (Figure S4). Surprisingly, quantitative analyses revealed there was no statistical difference in number of mitolysosomes between wild type and *Pink1* or *parkin* mutant animals for most cell types except for larval CNS where the difference in *Pink1* mutants just reached significance (Figures 3 and 4).

Finally, although mitolysosomes were not detectable in WT flight muscle we considered that since profound mitochondrial disruption occurs in flight muscle of *Pink1/parkin* mutants mitophagy may be induced as a homeostatic response. However, we again observed no mitophagy signal in *Pink1* or *parkin* mutants (data not shown). Thus, taken together these data indicate that loss of Pink1 or parkin does not substantially influence basal mitophagy in neuromuscular tissues.

The notion that PINK1 and parkin cooperate to mediate bulk degradation of mitochondria has become a dominant concept in the field of PD pathogenesis. While a large body of work has elucidated the molecular details of PINK1/parkin mediated stress-induced mitophagy *in vitro*, it has been an open question whether PINK1 and parkin perform equivalent functions under physiological conditions (Cummins and Gotz, 2017; Rodger et al., 2017; Whitworth and Pallanck, 2017). We have made use of two previously devised mitophagy reporter constructs to visualize mitophagy in fly models for the first time. Our observations reveal that basal mitophagy is highly prevalent in multiple *Drosophila* tissues, however, this mitophagic process is essentially unaffected by null mutations in *Pink1* or *parkin*.

The simplest interpretation of these results is that PINK1 and parkin do not in fact play an important role in basal mitophagy; however, this seems unlikely given the plethora of evidence supporting PINK1/parkin in some form of mitochondrial degradation. A number of alternative explanations can be proffered. One clear possibility is that another mode of mitophagy can be induced as a compensatory mechanism. Indeed, there is some precedence for this as it was recently reported that MUL1 acts redundantly alongside parkin (Rojansky et al., 2016; Yun et al., 2014). Moreover, while germline knockout of murine *PINK1* and *parkin* has been shown to have very mild phenotypes (reviewed in (Lee et al., 2012)), conditional post-natal loss of PINK1 or parkin has revealed robust loss of DA neurons (Lee et al., 2017; Shin et al., 2011). The authors propose that in germline knockouts an unknown mechanism is induced during development that compensated for loss of PINK1/parkin. One possibility that would fit this scenario is induction of an alternative mitophagy pathway or upregulation of other mitochondrial quality control processes. Clearly, more needs to be learned about the complex regulatory mechanisms governing mitochondrial turnover under physiological conditions.

Another obvious interpretation of the current findings is that basal mitophagy may reflect a distinct homeostatic mechanism from that in which PINK1/parkin function. Indeed, whether the observed basal mitophagy is operating as a quality control mechanism or some other homeostatic process, such as maintenance of mitochondrial levels, is unknown. As previously described, the majority of studies analyzing PINK1/parkin mitophagy have used exogenous stressors and parkin over-expression, leaving it unclear what, if any, physiological stimuli provoke the same PINK1/parkin response. Nevertheless, supporting the requirement for a ‘second hit’, DA neurons become selectively vulnerable to loss of parkin when mitochondrial DNA mutations are elevated (Pickrell et al., 2015).

Growing evidence indicates that PINK1/parkin-mediated turnover of mitochondrial components may occur in a piecemeal fashion. For instance, multiple studies have described the selective targeting of portions of mitochondria for turnover (Burman et al., 2017; Hamalainen et al., 2013; McLelland et al., 2016; McLelland et al., 2014; Yang and Yang, 2013), a process likely occurring via so called mitochondria-derived vesicles (MDVs) (Sugiura et al., 2014). That selective degradation of mitochondrial components may also occur *in vivo* is supported by compelling evidence using a mass spectrometric method to analyze isotope labeled mitochondrial protein turnover in flies (Vincow et al., 2013). In light of this, it is clear that further work is needed to define the conditions in which PINK1 and parkin promote mitochondrial homeostasis. Notably, it will be important to determine the nature of physiological stressors trigger PINK1/parkin mitochondrial turnover. The mitophagy reporter lines described here provide new model systems for studying physiological mitophagy and can be potentially used for screening for novel regulators. Greater insights into the physiological function of PINK1/parkin-mediated mitochondrial homeostasis will allow a clearer understanding of the pathogenic causes of PD, and thus a better prospect for more rational therapeutic interventions.

## Methods

### *Drosophila* stocks, husbandry and locomotor assay

Flies were raised under standard conditions at 25°C on food consisting of agar, cornmeal, molasses, propionic acid and yeast in a 12h:12h light:dark cycle. *Pink1*^*B9*^ mutants were a kind gift from Prof. J. Chung (Park et al., 2006), and the *park*^*25*^ mutants have been described previously (Greene et al., 2003). The following strains were obtained from the Bloomington *Drosophila* Stock Center (RRID:SCR_006457): *da-GAL4* (RRID:BDSC_55850), *nSyb-GAL4* (RRID:BDSC_68222), *Mef2-GAL4* (RRID:BDSC_27390), *Atg5*-RNAi (RRID:BDSC_34899), *TH-GAL4* (RRID:BDSC_8848). Locomotor (climbing) ability was assessed using a negative geotaxis assay as previously described (Greene et al., 2003). Transgenic expression was induced by *da-GAL4*, and adult male flies were tested 1-3 days after eclosion. Since X chromosome non-dysjunction is present in multiple balanced *Pink1*^*B9*^ mutant stocks, correct genotypes were determined by either combining paternal animals with X chromosome markers or by PCR-based genotyping of discarded tissue following dissection. For deferiprone treatment, mito-QC or mt-Keima expressing animals were raised on normal food dosed with deferiprone (LKT Laboratories, Cat# D1720) to a final concentration of 65 μM.

### Transgenic line construction

A plasmid containing the mito-QC sequence was obtained through the MRC Protein Phosphorylation and Ubiquitylation Unit Reagents and Services facility (MRC PPU, College of Life Sciences, University of Dundee, Scotland; mrcppureagents.dundee.ac.uk: pBabe.hygro-mCherry-GFP FIS 101-152(end) [DU40799]) (Allen et al., 2013). PCR primers containing *Eco*RI and *Xba*I restriction sites were used to amplify mito-QC, and following restriction digestion and purification by standard methods, was cloned into complementary sites in a pUAST.attB transgenesis vector. Constructs were verified by sequencing before sending for transgenesis by BestGene Inc. UAS-mito-QC was integrated into attP16 and attP2 sites, and verified by PCR. Several transformant lines were tested for consistency before selecting a single line of each integration site for further study.

mt-Keima (Katayama et al., 2011) was amplified and cloned from mt-Keima (h)-pIND(SP1) plasmid into a *Drosophila* expression vector pUASt. pUASt-mt-Keima construct was injected and recombined in *w*^*1118*^ fly embryos at BestGene Inc. Four independent UAS-mt-Keima strains were selected: UAS-mtKeima M7 (II), UAS-mt-Keima M3 (II), UAS-mt-Keima M2 (III) and UAS-mt-Keima M4 (III) based on detection of fluorescent mt-Keima signal in larval eyes by crossing them with GMR-GAL4. UAS-mt-Keima M2 flies were genetically recombined with Tub-GAL4/TM3 (III) flies to finally generate Tub-GAL4, UAS-mt-Keima M2/ TM3.

### Antibodies and dyes

The following antibodies and dyes were used: ATP5A (Abcam Cat# ab14748, RRID:AB_301447; 1:500), Actin (Millipore Cat# MAB1501, RRID:AB_2223041; 1:5000), GFP (Abcam Cat# ab290, RRID:AB_303395; 1:2000), DsRed (Clontech Laboratories, Inc. Cat# 632496, RRID:AB_10013483; 1:1000), Keima-Red (MBL International Cat# M126-3, RRID:AB_10210643; 1:1000), alpha-Tubulin (Sigma-Aldrich Cat# T6793, RRID:AB_477585; 1:5000), LysoTracker™ Deep Red (Invitrogen, L12492; 500nM). Secondary antibodies used, for immunoblotting: anti-rabbit HRP (Thermo Fisher Scientific Cat# G-21234, RRID:AB_2536530; 1:10,000), anti-Mouse HRP (Abcam Cat# ab67891, RRID:AB_1140250; 1:10,000); for immunofluorescence: anti-Mouse AF647 (Thermo Fisher Scientific Cat# A-21235, RRID:AB_2535804; 1:200).

### Immunoblotting

Protein samples isolated from whole adult fly were prepared in lysis buffer [50 mM Tris·HCl, 150 mM NaCl, 10% (vol/vol) glycerol, 1% Triton X-100, 10 mM N-ethylmaleimide, 2 mM EGTA, 1 mM, MgCl_2_, 50 μM MG-132, and protease inhibitor mixture (Roche)], resolved by SDS-PAGE using 12% precast gels (Bio-Rad) and transferred onto nitrocellulose membrane (Bio-Rad) using a Bio-Rad Transblot Transfer Apparatus. Membranes were blocked with 5% skimmed milk in TBST (Tris-buffered saline with 0.1% Tween-20) for 1 h at RT and probed with the appropriate primary antibodies diluted in the blocking solution overnight at 4°C. Membranes were washed repeatedly in TBST, then appropriate horse radish peroxidase-conjugated secondary antibodies (Dako) were incubated for 1 h at RT. Detection was achieved with ECL-Prime detection kit (Amersham). As a loading control, membranes were incubated with anti-alpha-Tubulin or anti-Actin antibodies for 1 h at RT.

### Immunohistochemistry and sample preparation

Mito-QC larvae, adult brains and thoracic muscles were dissected in PBS and fixed in 4% formaldehyde (pH 7.0) for 30 min. For immunostaining, larval tissues were permeabilized in 0.3% Triton X-100 for 30 min and blocked with 0.3% Triton X-100 and 1% goat serum in PBS for 1 h at RT. Tissues were incubated with ATP5A antibody diluted in 0.3% Triton X-100 and 1% goat serum in PBS overnight at 4°C, then rinsed 3 times 10 min with 0.3% Triton X-100 in PBS and incubated with the appropriate fluorescent secondary antibodies for 2 h at RT. The tissues were washed 2 times in PBS and mounted on slides using Prolong Diamond Antifade mounting medium (Thermo Fischer Scientific). Mt-Keima larvae were dissected and mounted in PBS. Tissues were rapidly imaged via sequential excitations (458 nm, green; 561 nm, red) being captured at 578 to 638 nm emission range.

### LysoTracker assay

Larvae were dissected in PBS. The epidermal cells were stained for 2 min with LysoTracker™ Deep Red (Invitrogen) diluted in PBS (final concentration of 500 nM). Mounting was performed in PBS and the samples were rapidly visualized by confocal microscopy.

### Microscopy

Fluorescence imaging was conducted using a Zeiss LSM 880 confocal microscope (Carl Zeiss MicroImaging) equipped with Nikon Plan-Apochromat 40x/1.3 NA and 63x/1.4 NA oil immersion objectives, or an Andor Dragonfly spinning disk confocal microscope (Andor) equipped with a Nikon Plan-Apochromat 100x/1.45 NA oil immersion objective. Z-stacks were acquired at 0.5 μm steps.

### Image analysis and quantification

Confocal images were processed using FIJI (ImageJ) software (RRID:SCR_002285) for figure presentation. For quantification of mitolysosomes, spinning disk microscopy generated images were processed using Imaris (version 9.0.2) analysis software (BitPlane) (RRID:SCR_007370) to identify and count individual ‘red-only’ puncta. The GFP and mCherry signals were adjusted to reduce background noise and retain only the distinct mitochondria network and red puncta, respectively. A surface rendered 3D structure corresponding to the mitochondria network was generated using the GFP signal. This volume was subtracted from the red channel to retain the mCherry signal that did not co-localize with the GFP-labelled mitochondria network. The mitolysosome puncta were selected according to their intensity and an estimated size of 0.5 μm diameter, previously measured with Imaris. Additionally, the puncta were filtered with a minimum size cutoff of 0.2 μm diameter. The remaining puncta were counted as the number of mitolysosomes.

Epidermal cells and larval CNS soma were analyzed individually where discrete cells could be distinguished. For DA neurons and adult CNS soma, small groups of cells were analyzed together and an average score calculated per cell.

The average number of mitolysosomes per cell was calculated per animal based on several images. Data points in the quantification charts show mean mitolysosomes per cell for individual animals, where n = 6 animals for all conditions except 10 days old DA neurons where n = 3 animals.

### Statistical analysis

For behavioral analyses, Kruskal-Wallis non-parametric test with Dunn’s post-hoc correction for multiple comparisons was used. Mitolysosomes were analyzed by Welch’s *t*-test. Analyses were performed using Prism 7 software.

## Acknowledgements

This work is supported by MRC core funding (MC-A070-5PSB0) and ERC Starting grant (DYNAMITO; 309742), and MRC studentship to J.J.L. Stocks were obtained from the Bloomington Drosophila Stock Center which is supported by grant NIH P40OD018537. We are grateful to Ian Ganley and Tom McWilliams for providing the mito-QC construct, discussions and communicating results prior to publication. mt-Keima (h)-pIND(SP1) plasmid was kindly provided by Hiroko Sakurai (RIKEN Brain Science Institute, Japan). We would also like to thanks Juanma Ramirez (UPV-EHU) for his help screening the insertion sites in the different UAS-mt-Keima founders.

## Author Contributions

M.J.C. and A.J.W. conceived the study. A.S-M., U.G., M.J.C. and A.J.W. designed experiments and supervised the work. J.J.L. and A.S-M. performed experiments. A.M.Z. generated UAS-mt-Keima lines. C.B. gave expert advice with microscopy and image analysis. A.J.W. wrote the manuscript with input from all authors.

## Supplementary Figure legends

**Figure S1. Expression of mito-QC and mt-Keima is benign in *Drosophila***. (A, B) Expression analysis of independent transgenic lines expressing (A) mito-QC and (B) mt-Keima induced by tubulin (tub) or daughterless (da) GAL4 lines. Immunoblots were probed with the indicated antibodies. Control genotype is *w*^*1118*^. (C-F) Locomotor assays for (C, D) climbing and (E, F) flight ability of the same genotypes analysed above. Charts show mean ± 95% CI. n>50 animals. Statistical analysis was determined by Kruskal-Wallis non-parametric test with Dunn’s post-hoc correction for multiple comparisons; ns = non-significant. Control genotypes for behavior is *da-GAL4/*+.

**Figure S2. Validation of mito-QC and mt-Keima reporter targeting**. Immunohistochemical and confocal imaging analysis of (A, C) larval epidermal cells and (B) adult flight muscle. (A, B) Native GFP fluorescence (green) and (C) 458 nm stimulated fluorescence (neutral pH) were compared to a mitochondrial protein distribution by immunostaining with anti-ATP5A (magenta). Confocal imaging analysis of larval epidermal cells for (A) mito-QC and (B) mt-Keima neutral and acidic pH spectra co-stained for LysoTracker to mark lysosomes. Fluorescence spectrum of neutral pH shown in green, acidic pH shown in red. LysoTracker is shown in white. Scale bars = A and C, top panels and B, 10 μm; A and C, bottom panels 4 μm. D and E = 10 μm. Genotypes analyzed were: (A, D) *da-GAL4/UAS-mito-QC*, (B) *Mef2-GAL4/UAS-mito-QC*, and (C, E) *tub-GAL4*, *UAS-mt-Keima*.

**Figure S3. Mitolysosome signal is lost upon autophagy inhibition and increased by deferiprone**. Confocal imaging analysis of larval epidermal cells for mito-QC and mt-Keima upon (A) expression of an *Atg5*-RNAi transgene to inhibit autophagy, or (B) exposure to the iron chelator deperiprone (65 μM) to induce mitophagy. Scale bars = 10 μm. Genotypes analyzed were: *da-GAL4/UAS-mito-QC*, and *tub-GAL4*, *UAS-mt-Keima*, plus UAS-*Atg5*-RNAi for panel A.

**Figure S4. Mitolysosome quantification workflow (mito-QC only)**. Z stack images were acquired with spinning disk for quantification. Where possible individual cells were isolated for workflow analysis. The contrast of the mitochondria network was adjusted to reduce the background, and a rendered 3D surface corresponding to the mitochondria network was generated. This surface was then subtracted to the mCherry channel to discard the mCherry signal overlapping the GFP-labelled mitochondria network leaving the red-only puncta. Genotypes shown is *da-GAL4/UAS-mito-QC*.

